# Genome-wide impact of codon usage bias on translation optimization in *Drosophila melanogaster*

**DOI:** 10.1101/2023.10.05.561139

**Authors:** Xinkai Wu, Jian-Rong Yang, Jian Lu

## Abstract

Accuracy and efficiency are fundamental characteristics of the translational process. Codon usage bias is widespread across species. Despite the long-standing association between codon optimization and improved translation, our understanding of the evolutionary basis and functional effects of codon optimization is limited. *Drosophila* has been widely used to study codon usage bias, but genome-scale experimental data on codon optimization and translation are scarce. We examined high-resolution mass spectrometry datasets from *D. melanogaster* development and employed different strategies to reduce bias when comparing translation error rates. We demonstrated that optimal codons have lower translation errors than nonoptimal codons after accounting for these biases. Our findings also shed light on codon-anticodon mismatches in translation errors. Through genomic-scale analysis of ribosome profiling data, we showed that optimal codons are translated more rapidly than nonoptimal codons in *D. melanogaster*. While we did not find conclusive evidence that natural selection favored synonymous mutations during the long-term evolution of the *D. melanogaste*r lineage after its divergence from *D. simulans*, we did find that positive selection drives codon optimization-related mutations in the *D. melanogaster* population. This study expands our understanding of the functional consequences of codon optimization, and serves as a foundation for future investigations into the molecular mechanisms governing gene expression evolution at the translation level.

## Introduction

The translation of messenger RNA (mRNA) into proteins is a fundamental process in all domains of life. However, translation is not completely error-free. A common type of translational error is amino acid misincorporation, occurring at a rate of approximately 10^−3^ to 10^−4^ per codon site ^1,2^. Despite their contribution to phenotypic plasticity ^3-5^, translation errors are generally detrimental to organisms as they reduce the functional activities of proteins and can lead to toxic protein misfolding and misinteraction ^6-12^. Consequently, it is not surprising that translational fidelity has been shown to be optimized through strong natural selection ^13-20^.

The optimization of translational fidelity, however, comes with its costs. Early studies have demonstrated a trade-off between accuracy and efficiency in mRNA translation ^21-23^. Increasing fidelity at the expense of efficiency may not necessarily be advantageous, as maintaining translational efficiency is crucial for ensuring the availability of limited resources like ribosomes and tRNAs. This is especially significant for rapidly dividing cells ^24,25^. Notably, when comparing the translational efficiency of the same codon in different genomic contexts, an increase in accuracy at the cost of efficiency can only be observed at genomic sites where amino acid misincorporations are highly detrimental to the cell ^26^. Mechanistically, the trade-off between accuracy and efficiency is underpinned by factors such as Magnesium concentration ^23^ and mRNA secondary structure ^26^.

However, beyond the aforementioned site-specific regulation, the orchestration of translational accuracy and efficiency encompasses a multitude of mechanisms, creating a more intricate relationship when viewed from various angles. For instance, the interplay between accuracy and efficiency across distinct types of codons is substantially influenced by tRNA selection during translational elongation ^15^. Firstly, codons associated with more abundant aminoacyl-tRNAs (aa-tRNAs) are translated more efficiently than codons associated with less abundant aa-tRNAs ^17,27,28^. Secondly, during translation elongation, a cognate aa-tRNA is favored for peptidyl transfer, whereas a noncognate aa-tRNA tends to dissociate from the ribosome in a proofreading step before peptidyl transfer ^29,30^. This raises the question of whether the accuracy-efficiency trade-off, which is observed among different genomic sites of the same type of codon, also applies to the accuracy-efficiency relationship across different codon types. To answer this question, comprehensive empirical data that provide matched estimations of accuracy and efficiency on a genome-wide scale are required. Recent technological advancements have facilitated this pursuit. On the one hand, ribosome profiling (Ribo-Seq) has emerged as a powerful tool for quantifying translational elongation rates by examining ribosome aggregation at specific genomic positions ^31-35^. On the other hand, advances in mass spectrometry analytical pipelines have enabled the systematic identification of translation errors at the proteome level, achieved by disentangling erroneously translated peptides from their correctly translated counterparts ^20,36^.

Apart from its functional genomic implications, the earlier mentioned question carries significant evolutionary implications, particularly in terms of the molecular mechanisms that underlie natural selection for synonymous codon usage bias (CUB). CUB refers to the phenomenon observed among codons that encode the same amino acid, where certain optimal or preferred codons are more frequently used than nonoptimal or rare codons. CUB is prevalent across organisms ^37^ and is influenced by mutation ^38-41^, drift ^42^, selection ^12,37,43-45^, or their combinations ^46,47^. Previous studies based on simple organisms such as bacteria and yeasts have shown that CUB significantly impacts the efficiency and accuracy of mRNA translation, potentially contributing to organismal fitness and adaptation ^15,16,18,19,44,48-52^. Despite this, the interplay between translation efficiency and accuracy at the individual codon level still poses a challenging aspect to fully understand and characterize, particularly when considering complex multicellular organisms.

*Drosophila* has been extensively utilized as a model to investigate CUB, predominantly in the realms of population genetics and evolutionary biology, with the underlying assumption of a link between CUB and translational optimization (e.g.,^53-55^). For instance, prior studies have demonstrated that evolutionarily conserved amino acid residues tend to favor preferred codons, implying that conserved residues necessitate heightened translation accuracy in *Drosophila* ^14^. Notably, preferred codons in *Drosophila* frequently terminate with G or C, possibly due to selection for codon bias ^56^. Synonymous sites generally experience weak selective constraints in *Drosophila* and other species ^57,58^. Analyzing fixed versus polymorphic synonymous mutations in five genes of *D. melanogaster* and *D. simulans* has revealed a preference for mutations from nonoptimal to optimal codons, while the reverse is disfavored in the lineage leading to *D. simulans* ^13^. However, this analysis has also indicated a reduction in selection pressure on synonymous mutations in the lineage of *D. melanogaster* after its divergence from the most recent common ancestor with *D. simulans*. The observed difference in these two lineages likely arose due to a smaller effective population size of *D. melanogaster* compared to *D. simulans* ^13,59,60^, a change in the selective coefficient on the synonymous sites due to changing ecological conditions ^61^, a shift in mutational bias towards A/T alleles in the *D. melanogaster* lineage ^62-65^, or a combination of these factors.

While theoretical studies have long suggested that natural selection for translation optimization leads to increased codon bias levels ^45,66^, experimental evidence confirming the impact of codon optimization on translation accuracy or speed in *Drosophila* remains limited. Notable experiments have shown that introducing less preferred synonymous codons into the *Adh* gene reduces ADH protein levels and ethanol tolerance in *D. melanogaster* ^67,68^. Additionally, in vitro translation experiments employing luciferase reporter genes have underscored codon optimization’s ability to enhance ribosome translation elongation rates in *D. melanogaster* ^69^. Despite these individual experiments, a notable gap persists in our understanding due to the absence of comprehensive genome-scale experimental data exploring the influence of codon optimization on translation in *Drosophila*. This gap is particularly striking, given the widespread use of *Drosophila* in CUB studies. Therefore, there is a pressing need for further research to fill this crucial knowledge gap comprehensively.

Significantly, a discrepancy arises concerning the existence of codon usage bias to codon optimization in *D. melanogaster*. On the one hand, prior studies focusing on a limited number of genes found no evidence of codon optimization driven by natural selection in the lineage leading to *D. melanogaster* ^13^. On the other hand, it is widely accepted that natural selection for translation optimization leads to higher codon bias levels ^45,66^. As a result, a crucial gap remains in connecting functional outcomes with the evolutionary drivers of CUB in this model organism.

This study aimed to address these questions by evaluating translational efficiency and accuracy at the genome-wide level in *D. melanogaster*. We analyzed high-resolution mass spectrometry datasets ^70^ to measure translation errors and utilized Ribo-Seq data that we previously generated ^71^ to assess elongation rates at the codon level in *D. melanogaster*. Our findings provide genome-wide evidence that codon optimization increases both translation accuracy and efficiency in this organism. Additionally, we revisited the evolutionary forces influencing the evolution of synonymous mutations using population genomic data from *D. melanogaster*. Our study serves to bridge an existing knowledge gap and contribute to a more comprehensive understanding of how natural selection influences codon usage in the translational process during evolution.

## Results

### Detection of amino acid misincorporation using mass spectrometry data

Either tRNA mischarging or codon-anticodon mispairing causes amino acid misincorporation, leading to translation errors (Fig. 1a). To detect and analyze individual misincorporation events, we thoroughly examined high-resolution mass spectrometry datasets comprising 17 different samples collected from 15 representative time points during *D. melanogaster* development ^70^. Each of these samples consisted of four biological replicates (see *Methods*). Since the mass spectrometry data were generated using the Oregon-R strain ^70^, our initial step involved generating the Oregon-R reference coding sequences by incorporating single nucleotide polymorphisms (SNPs) into the reference genome based on the ISO-1 strain of *D. melanogaster* (Supplementary Fig. 1) ^72^. On the basis of these reference sequences of the Oregon-R strain, we used MaxQuant ^73^ to distinguish between base peptides (those without errors or amino acid substitutions) and dependent peptides (those with residual modifications or substitutions). Following a method previously described ^20^, we identified amino acid changes by comparing mass differences between dependent and base peptides, and inferred possible mispairings between codons and anticodons (Fig. 1b). Given the rarity of translation errors, we considered amino acid substitutions present in no more than three of the 68 samples (four independent replicates for each of the 17 distinct samples) to be translation errors (Supplementary Fig. 2). Additionally, we excluded known adenosine-to-inosine (A-to-I) RNA editing sites in *D. melanogaster* ^74^, as A-to-I RNA editing can post-transcriptionally alter the protein sequence ^75,76^. After applying these criteria, we successfully identified translation error events in 13 to 61 genomic sites in each sample (Fig. 1c). Overall, we discovered a total of 1,374 distinct genomic sites with translation errors (Supplementary Fig. 3), out of which 992 were found in one sample, 265 in two samples, and 117 in three samples (Supplementary Fig. 2).

**Fig. 1.**
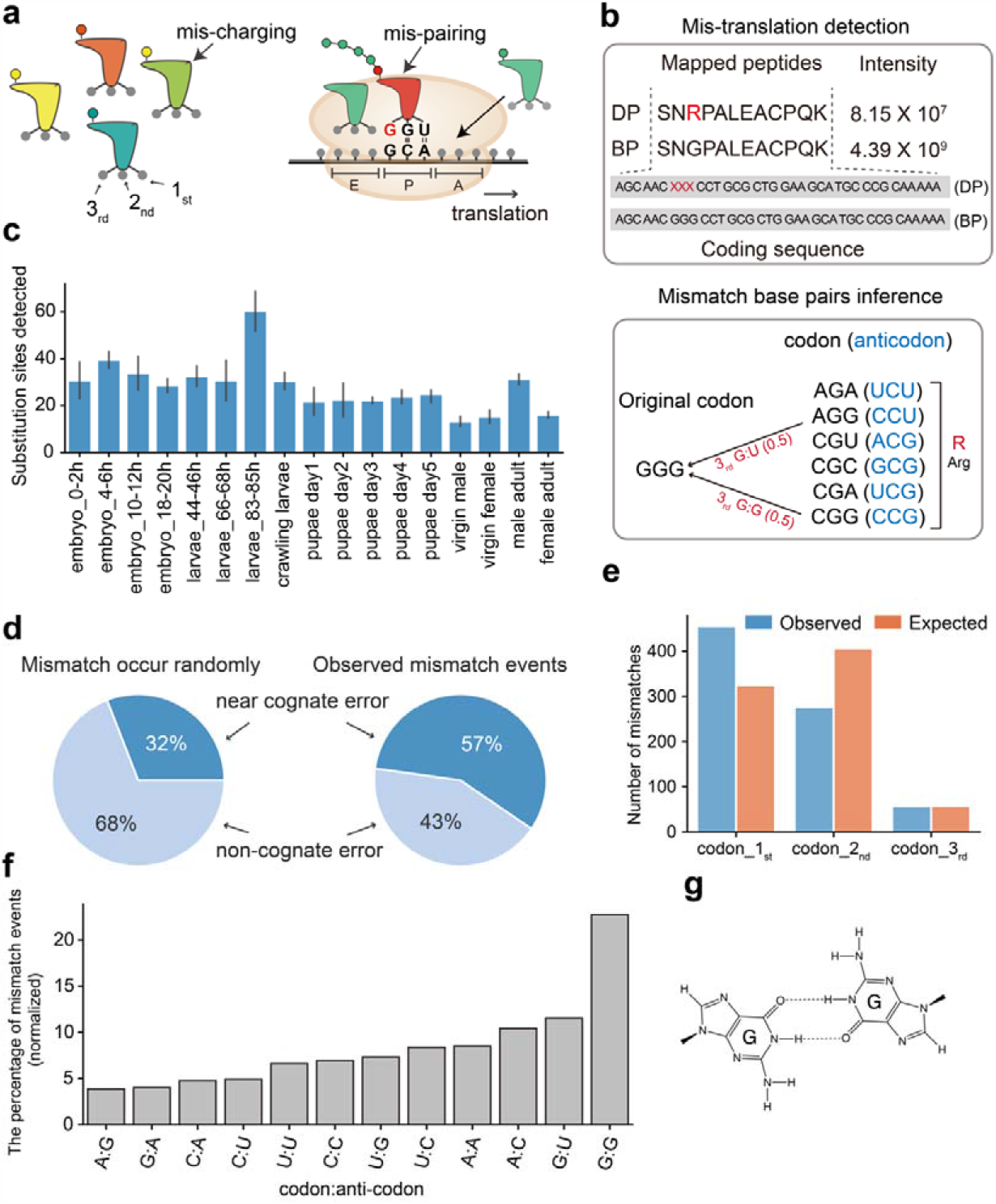
Summary of translation errors. **a**. Causes of Translation Errors: tRNA Mischarging and Codon-Anticodon Mispairing. **b**. Illustration of an identified translation error. Upper panel: the error events can be inferred by comparing the mass difference between dependent peptide (DP) and base peptide (BP); Lower panel: the possible mispairing between a codon and anticodon can be inferred by comparing the amino acids in the DP versus BP. **c**. The number of genomic sites that showed translation errors in the samples. **d**. The expected fractions of NeCE and NoCE occurrences under randomness (left) and the observed fractions of NeCE and NoCE occurrences. **e**. The number of NeCE mismatches (*y*-axis) at each position of codons (the *x*-axis). **f**. The percentages (*y*-axis) of base pairs mismatched (*x*-axis) in the codon-anticodon pairing in NeCE occurrences. **g**. G:G mismatch can form two stable hydrogen bonds.

As observed in *E. coli*, we also found that near-cognate errors (NeCEs) caused by one codon-anticodon mismatch occur more frequently than noncognate errors (NoCEs) resulting from multiple mismatches (Fig. 1d). Among the 1,374 translation error instances identified, 57% (788) were NeCEs, and 43% (586) were NoCEs, significantly deviating from the expected 32% NeCE and 68% NoCE under the null hypothesis that a codon has equal probability to be translated into each of the 19 other amino acids (see *Methods*). Upon comparing mismatch distributions in NeCE occurrences across codon-anticodon pair sites, we found similar occurrences of mismatches in the third codon position in the observed versus simulated random data (Fig. 1e; *Methods*), supporting the concept of wobble base pairing ^77^. However, the observed NeCE mismatches were significantly enriched in the first position but underrepresented in the second codon position (*P* < 0.001, χ^*2*^ test; Fig. 1e). These data suggest that mismatches in the first position of the codon significantly contribute to translation errors. Our finding also aligns with earlier structural insights that ribosomal decoding is largely determined by the stability of the base pair at the second position of the codon-anticodon interaction ^78^.

Notably, G:G mismatches were the most prevalent, constituting 25% of NeCE instances (198/788) and 23% of all mismatch events after normalizing by codon base compositions (Fig. 1f). The presence of two stable hydrogen bonds in G:G pairing^79^ suggests its high tolerance as a codon-anticodon mismatch in translation elongation (Fig. 1g), making it a significant source of translation errors.

### Optimal codons are translated more accurately in *D. melanogaster*

The translation error rates have been shown to vary among codons ^20,52,80,81^. To test whether optimal codons are translated more accurately in *Drosophila*, we calculated the relative synonymous codon usage (RSCU) ^82^ values for each specific codon. Unlike the traditional RSCU process that uses a small number of highly expressed genes to calculate codon usage frequency, we took into account the gene expression levels from various developmental stages of *D. melanogaster* in our analysis, which allowed us to determine a more representative average usage frequency for codons in the transcriptomes (*Methods*). The codons with RSCU ≥ 1 were classified as optimal, whereas the rest were considered nonoptimal (Supplementary Data 1). For each of the 61 codon types, we quantitatively estimated the translation error by comparing the signal intensity of the dependent peptides containing that codon to the signal intensity of base and dependent peptides containing that codon type in a sample, while adjusting for codon compositions in the mass spectrometry data of the given sample (Fig. 1b and Supplementary Data 2). In other words, our translation error estimation for each codon type was equivalent to the fraction of misincorporated amino acids out of the total number of amino acids encoded by that codon in the proteome.

One potential bias in investigating the impact of codon optimization on translation error is that, due to the low translation error rate (10^−3^ to 10^−4^ per codon site ^1,2^), sampling bias might cause some rare codons not displaying any detected translation errors in the mass spectrometry data. To illustrate, we noticed that when not accounting for the bias from codon usage frequency in the analysis, optimal codons, which are typically more frequently used, showed more translation errors than nonoptimal ones, which are less frequently used (Wilcoxon rank-sum test, *P* = 4.3×10□^13^, Fig. 2a). To address this, we simulated error detection across all codon types using a uniform error rate of 2.37×10□□ errors per amino acid. This rate was determined by attributing a zero error rate to codons with no detected error events in a library and computing the mean errors for all codon types (*Methods*). As anticipated, many rare codons did not exhibit translation errors when their corresponding peptides have only low signal intensity in the mass spectrum. However, increasing throughput of mass spectrometry led to more error detections (Fig. 2b). Moreover, the simulations revealed that translation errors were more likely to be detected for more frequently used codons (Fig. 2c). In conclusion, our simulations suggest that the infrequent usage of rare or nonoptimal codons, along with low translation error rates, can significantly bias translation error rate estimation. Therefore, caution should be exercised when comparing translation error rates between codons, particularly when codon usage varies considerably.

**Fig. 2.**
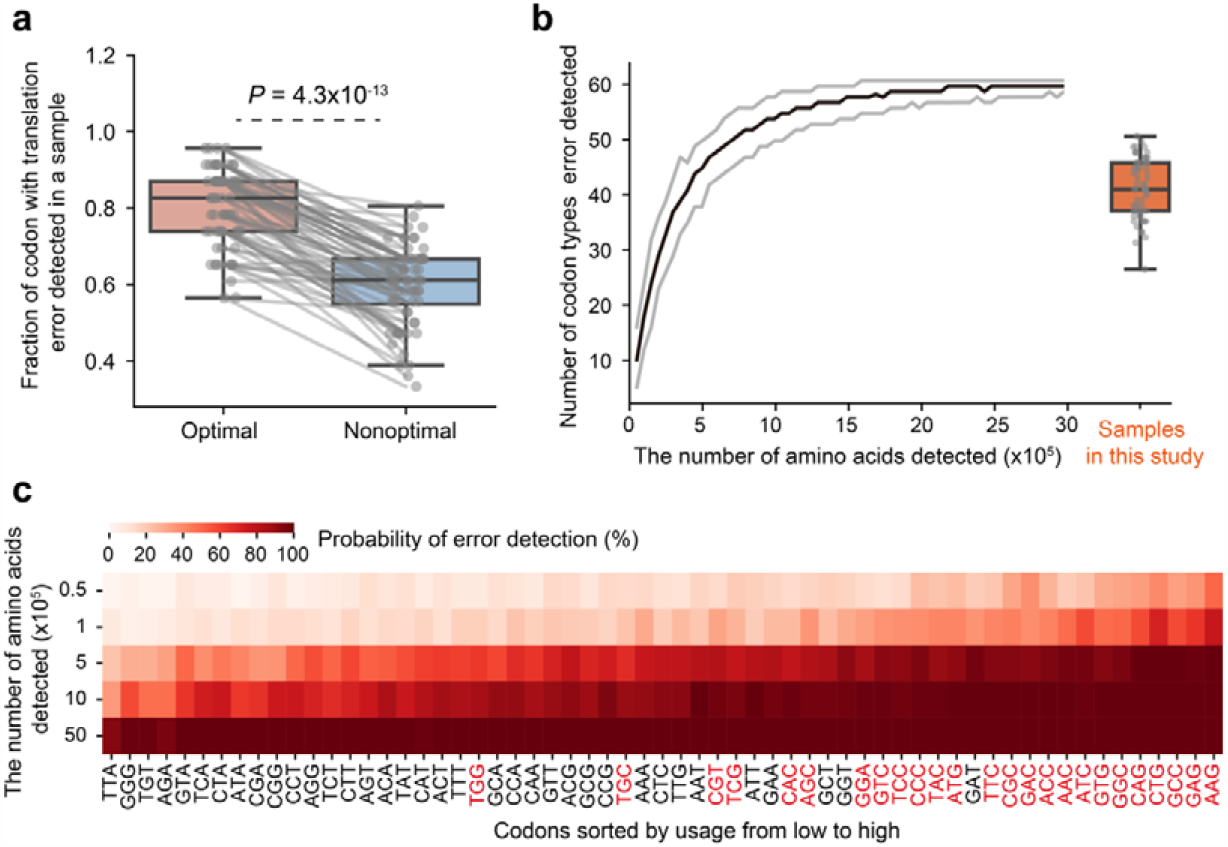
The influence of mass spectrometry throughput on error identification. **a**. Fraction (*y*-axis) of optimal codons and nonoptimal codons identified with translation error events in a mass spectrometry library. Each dot represents the proportion of optimal codons or nonoptimal codons in a sample with at least one translation error. Points in the same sample are connected by a solid gray line. The box plot shows the center line (median), box limits (upper and lower quartiles), and whiskers (1.5 times the interquartile range). **b**. The number of identified codon types carrying translation errors (*y*-axis) under different mass spectrometry throughputs (*x*-axis). Each simulation was independently repeated 100 times. Median values are in black, and 95% confidence intervals are shown in gray. The orange boxplot represents the distribution of identified codon types carrying translation errors in the 68 samples in this study. **c**. Probability of identification of translation error events for a codon at different mass spectrometry throughputs (*y*-axis). The color in each square represents the percentage of times that type of codon was mistranslated at a certain throughput in 100 independent simulations. Optimal codons are highlighted in red in *x*-axis.

We employed three strategies to reduce bias in comparing translation error rates. First, in each sample, we considered codons without detected errors to have a 0 error rate, grouped nonoptimal (or optimal) codons by the amino acid they encoded, and calculated the mean error rate of the nonoptimal (or optimal) codons for that amino acid. This method increased the sample size of nonoptimal (or optimal) codons, thereby reducing the effect of biased error rate estimation. Subsequently, we performed a linear mixed model analysis to compare the difference in translation error rates between optimal and nonoptimal codons, with developmental time points, amino acid types, and codon category (optimal or nonoptimal) as fixed effects, and experimental replicates as random effects (*Methods*). The linear mixed regression analysis demonstrated that optimal codons exhibited significantly lower translation error rates than nonoptimal codons (*P* = 0.031).

In the second strategy, we omitted codons without error events from each library, resulting in an average error rate of (3.50 ± 0.44)×10^−4^ errors per amino acid, which falls within the range of previously estimated error rates in bacteria (between 10^−4^ and 10^−3^ errors per amino acid)^1,2^. We also employed a linear mixed model to analyze the relationship between codon translation error rates and RSCU, with the experimental replicates as random effects. The results revealed a significant negative correlation between the translation error rate (log_10_) and the RSCU value (the slope coefficient = -0.141, *P* < 0.001, Fig. 3a), reinforcing the notion that optimal codons exhibit lower translation error rates than nonoptimal codons.

**Fig. 3.**
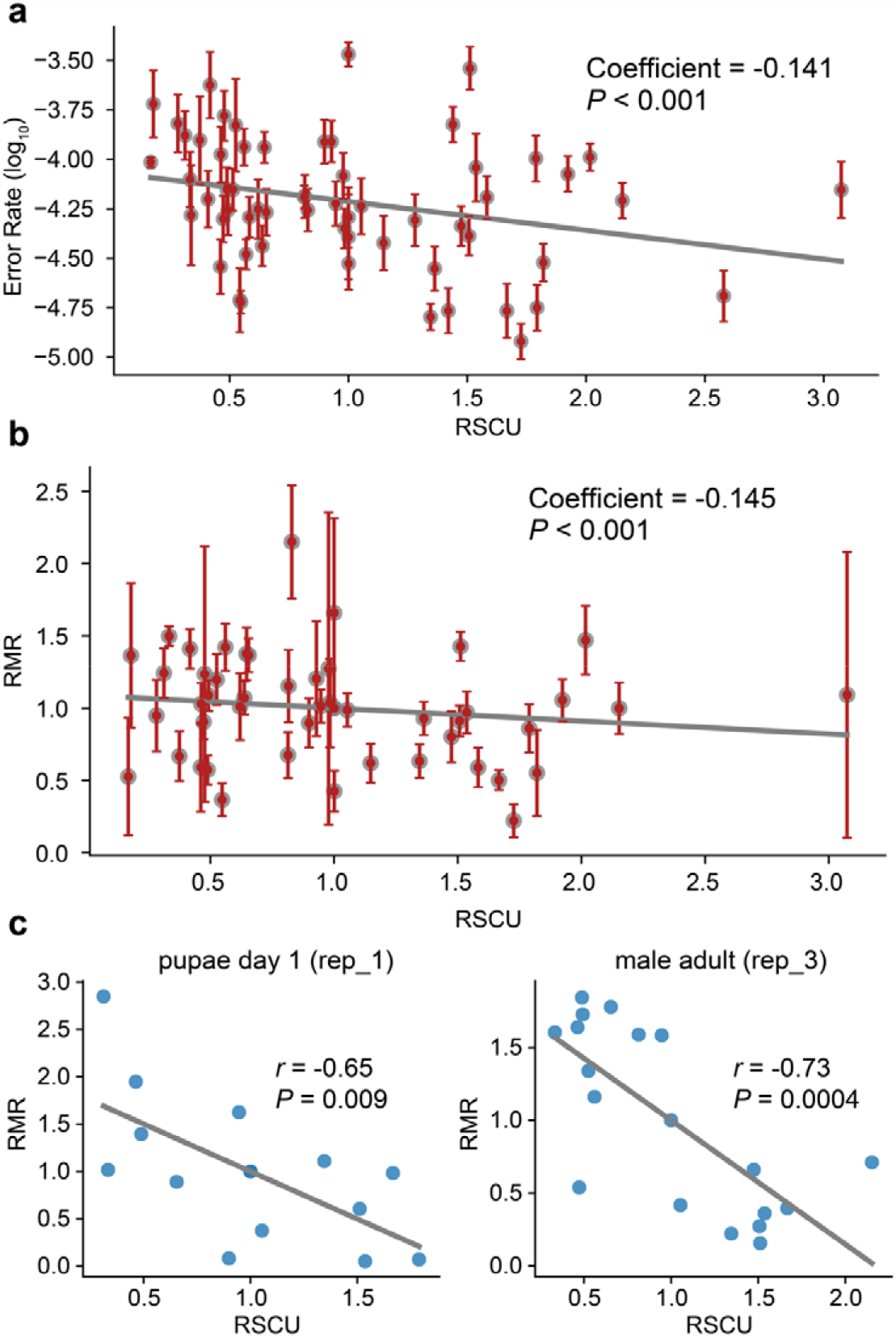
Relationship between translation error rate and codon optimality. **a**. An inverse relationship between the translation error rate (*y*-axis) and the RSCU value (*x*-axis) of a codon. In each sample, codons without translation errors were excluded. Each point represents a codon, with the *y*-axis representing the average error rate of that codon and error bars representing the standard errors of the translation error rate across different samples. The coefficient of the linear mixed model and *P* value are indicated. **b**. Correlation between relative mistranslation rate (RMR) (*y*-axis) and RSCU (*x*-axis). Only amino acids with all the synonymous codons identified as having translation errors in each sample were considered in the analysis. Each point represents the average relative error rate identified in different samples, with error bars indicating the standard error. The solid line represents the linear regression fitted line. The coefficient of the linear mixed model and the *P* value are shown in the plot. **c**. Two specific examples from (**b**) showing a significant negative correlation between RSCU and RMR.

In the third method, following the approach of a previous study ^52^, we considered only amino acids where all synonymous codons showed translation errors in a sample and calculated the relative translation error rate (RMR) by dividing a codon’s error rate by the average error rate of its synonymous codons. We found a significant negative correlation between RMR and RSCU when pooling data of all samples together (the slope coefficient = -0.145, *P* < 0.001, Fig. 3b). Fig. 3c provides two examples, one from pupae and the other from male adults.

Collectively, these findings suggest that although the estimation of translation error rates is often influenced by biases in codon occurrences, optimal codons generally exhibit lower translation errors than nonoptimal codons after accounting for these biases.

### Optimal codons are translated faster than nonoptimal codons in *D. melanogaster*

Introducing unpreferred synonymous codons into the *Adh* gene into *D. melanogaster* reduces ADH protein levels ^67,68^. In vitro translation experiments with luciferase reporter genes have shown that codon optimization increases the translation elongation rate in *D. melanogaster* ^69^. Given the variable codon usage bias among genes, we investigated whether this pattern could be detected using Ribo-Seq data reflecting in vivo conditions. To measure the impact of codon optimization on translation elongation, we analyzed ribosome profiling data that we had previously collected across various developmental stages and tissues of *D. melanogaster* ^71^, enabling us to calculate the local elongation rate of the ribosome. This dataset included ten samples, encompassing 0–2h, 2–6h, 6–12h, and 12–24h embryos; larvae, pupae, female and male bodies; and female and male heads. We mapped the ribosome-protected fragments (RPFs) to the reference genome, inferred the location of the A-site on each RPF, and calculated the ribosome density on each codon position in the coding region (CDS) of a gene (*Methods*). To account for differences in translation initiation rates between genes, we normalized the ribosome density on each codon position in a CDS to the average ribosome density of codons within that CDS. Overall, our analytical procedure aligns with the RibosOme Stalling Estimator (ROSE) framework, which calculates the likelihood of a ribosome stalling event at each genomic position ^34^. We excluded the first 15 and last five codons of a CDS from the analysis, as they are known to produce aberrant ribosome blockage due to translation start or termination. We also removed genes with insufficient RPF coverage (RPF coverage < 50%) or short CDSs (< 150 codons).

As Ribo-Seq can only capture a stationary snapshot of the translation processes, it is unsurprising that 20–33% of the codons exhibited no RPF detection in the expressed genes in a sample (Supplementary Fig. 4). These zero-RPF coverage codons could be under-sequenced, untranslated, or rapidly translated. Here, we only focused on the codons evidenced of translation, showing at least one RPF coverage. Optimal codons displayed significantly lower normalized ribosome density than nonoptimal codons (Fig. 4a and Supplementary Fig. 5), suggesting faster translation. We hypothesized that optimal codons would be substantially enriched in the bottom 0.1% of codons exhibiting the lowest normalized ribosome density (excluding codons without RPF coverage; see Fig. 4b for 0–2h embryos). Indeed, in all ten samples, optimal codons were significantly enriched among these rapidly elongating codons (Fig. 4c). To assess whether nonoptimal codons were enriched at stalling sites, we selected the top 0.1% of codons with the highest normalized ribosome densities as stalling sites within a sample. In nine out of ten samples, the nonoptimal codons were significantly enriched in these slowly elongating codons (Fig. 4d). Collectively, our data provide genome-wide evidence that in *Drosophila* tissues, optimal codons are translated more rapidly than their nonoptimal counterparts.

**Fig. 4.**
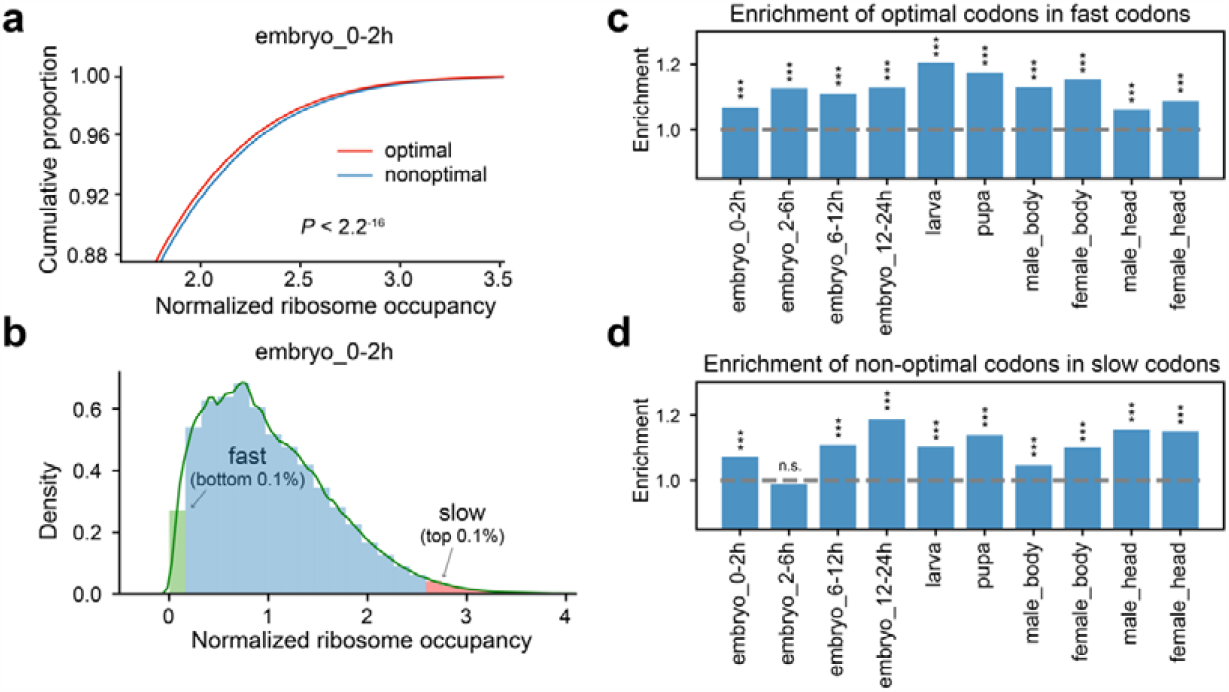
Impact of codon optimization on ribosome elongation rates. **a**. Comparison of normalized ribosome densities between optimal and nonoptimal codons in 0–2h embryos. The *P* value was calculated using the Wilcoxon rank-sum test. **b**. The distribution of log2-transformed normalized A-site occupancy in 0–2h embryos (zero-RPF covered codons were excluded). **c**. Enrichment of optimal codons in the bottom 0.1% codons showing the lowest normalized RPF coverage. **d**. Enrichment of nonoptimal codons in the top 0.1% of codons showing the highest normalized RPF coverage. ***, *P* < 0.001; n.s., not significant. The χ^*2*^ test was used in (**c**) and (**d**) to compute the *P* value in each sample.

### Detecting positive selection on the synonymous mutations that change codon optimality in *D. melanogaster*

To detect the signature of positive selection on synonymous changes in the lineage of *D. melanogaster* at the genome-wide level, we carried out the asymptotic McDonald-Kreitman test ^83,84^. As a neutral baseline for comparison, we focused on the 8–30 nt region of short introns (≤ 65 nt), which are considered neutral in *Drosophila* ^85-87^ (*Methods*). We used *D. yakuba* as an outgroup to polarize the fixed synonymous mutations in *D. melanogaster* after it diverged from *D. simulans* approximately 5.4 million years ago ^88^. We employed the polymorphic data from the *Drosophila* Genetic Reference Panel 2 (DGRP2) database of *D. melanogaster* ^89,90^ in the asymptotic MK test. We categorized fixed and polymorphic synonymous mutations into two groups: optimization (increasing RSCU) and deoptimization (decreasing RSCU), estimating the fraction of synonymous substitutions driven to fixation by positive selection (α) for each mutation category. Our findings revealed an α of -0.025 (95% confidence interval (CI): -0.076 to 0.026) for codon optimization substitutions and 0.014 (95% CI: -0.039 to 0.066) for codon deoptimization substitutions (Fig. 5a and 5b). These results corroborated previous observations based on a few genes, indicating the absence of evidence for positive selection on the synonymous mutations that became fixed in the lineage of *D. melanogaster* ^13^.

**Fig. 5.**
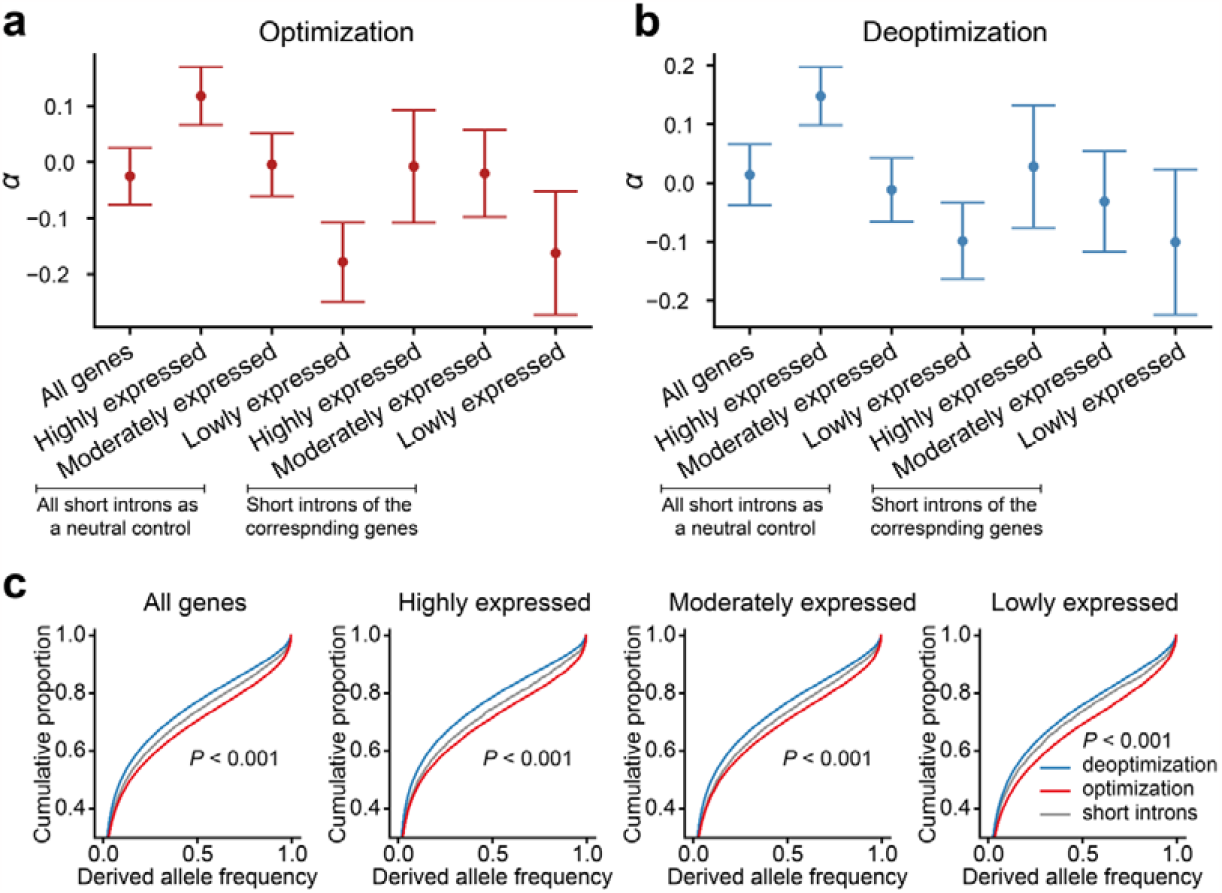
Positive selection of synonymous mutations altering codon usage frequency in *Drosophila*. **a-b**. Proportions (α) of positively selected synonymous mutations that have become fixed in the *D. melanogaster* lineage after diverging from *D. simulans*. The plot displays the median values (points) and their corresponding 95% confidence intervals (error bars) for α. Genes were categorized into three groups based on their median expression levels. It is important to note that using total short introns as a neutral control will yield different α values compared to using short introns from genes within corresponding expression categories. **c**. Derived allele frequencies (DAF) of synonymous mutations associated with codon optimization or deoptimization in the DGRP2 *D. melanogaster* data. Codon-optimizing mutations exhibited significantly higher DAF than neutral controls (mutations on short introns) in genes with different expression levels. Conversely, codon-deoptimizing mutations showed significantly lower DAF compared to neutral controls. The *P* value was calculated using the Wilcoxon rank-sum test.

As expected ^91-96^, when categorizing *D. melanogaster* genes into three groups based on their median expression levels (see *Methods*), we observed substantial differences in the prevalence of optimal codons. The proportions of optimal codons within the highly (*n*=4,565 genes), moderately (*n*=4,817), and lowly (*n*=4,517) expressed gene categories were 0.607, 0.556, and 0.523, respectively (Supplementary Fig. 6). For highly expressed genes, our results were somewhat equivocal. When we employed all short introns as a neutral control, we observed α values of 0.118 (0.066 to 0.170) for codon optimization and 0.148 (0.098 to 0.198) for codon deoptimization. However, when using only the short introns from these highly expressed genes as a neutral control, α values were -0.007 (−0.108 to 0.093) for codon optimization and 0.028 (−0.077 to 0.133) for codon deoptimization (Fig. 5a and 5b). Conversely, we found no positive selection on synonymous mutations in moderately and lowly expressed genes (Fig. 5a and 5b). Taken together, we did not find conclusive evidence that natural selection has favored synonymous mutations during the long-term evolution of the *D. melanogaster* lineage following its divergence from *D. simulans*.

The absence of positive selection on synonymous mutations in the *D. melanogaster* lineage could be attributed to various factors, including a smaller effective population size in *D. melanogaster* compared to *D. simulans* ^13,59,60^, changes in selection intensity due to shifting ecological conditions ^61^, and a stronger mutational bias from C/G towards A/T alleles in the *D. melanogaster* lineage ^62-65^. Temporal fluctuations or declines in these factors, coupled with their intricate interactions, could have potentially masked the signals of positive selection on synonymous mutations.

In contrast, a comparison of different classes of synonymous mutations in the *D. melanogaster* population may offer an alternative approach to detect signals of positive selection. This approach is less susceptible to factors that could have fluctuated or continued to evolve during long-term evolution. However, conflicting results have been obtained from previous studies on this issue. While some studies provided evidence for selection favoring preferred codons ^57,59,97^, others did not find support for this conclusion ^61,98,99^. To address this discrepancy, we analyzed the frequencies of codon-optimizing and deoptimizing mutations using the larger sample size provided by the DGRP2 dataset. Our results revealed that codon optimization mutations exhibit significantly higher derived allele synonymous frequencies compared to codon deoptimization mutations (*P* < 0.001, Wilcoxon rank-sum test; Fig. 5c). Furthermore, codon-optimizing mutations showed significantly higher derived allele frequencies compared to mutations within short introns (*P* < 0.001), while codon-deoptimizing mutations showed significantly lower derived allele frequencies than the mutations within short introns (*P* < 0.001). These trends persisted when analyzing genes based on different expression categories (Fig. 5c). In summary, these findings support the hypothesis that codon-optimizing mutations are favored by natural selection, while codon-deoptimization mutations are subject to negative selection in *D. melanogaster* populations.

## Discussion

Both translational accuracy and speed are crucial for efficient cellular protein synthesis. Decades of CUB studies in *Drosophila*, mainly from the angles of population genetics and evolutionary biology (e.g., ^13,14,45,53,54,100,101^), have bolstered the notion that natural selection drives biased synonymous codon usage to enhance protein synthesis accuracy and/or speed ^14,45,66^. For instance, it was previously shown that preferred codons were significantly more prevalent at conserved amino acid codons than at non-conserved ones in 38 genes between *D. melanogaster* and other species ^14^. This pattern was also evident when comparing the genomes of *D. melanogaster* with closely related species, *D. simulans* and *D. yakuba* (see Supplementary Fig. 7). Although these findings support the notion that conserved amino acid residues necessitate more accurate translation by using optimal codons ^14^, there is a noticeable gap in empirical data directly validating the hypothesis that codon optimization boosts protein translation accuracy. Similarly, few pieces of evidence confirm that codon optimization increases translation elongation speed in *Drosophila*, except for some case studies, like the *Adh* gene ^67,68^, and in vitro translation experiments using the luciferase reporter gene ^69^. In this genome-wide study, we offer direct evidence that codon optimization enhances translation accuracy and elongation speed in *D. melanogaster*. Our population genomics data analysis further underscores the concept that natural selection prefers codon-optimizing mutations and negatively selects against codon-deoptimizing mutations in the population of *D. melanogaster*. These insights serve as a bridge, linking functional results with the evolutionary underpinnings of CUB in this model species.

This study identified translational errors by analyzing high-resolution mass spectrometry datasets containing samples from 15 developmental time points in *D. melanogaster* ^70^. While past research determined error rates by contrasting signal intensities of correct and incorrect peptides^20^, this method might skew the error rate estimation since it overlooks translation events at error-free sites. To address this, we evaluated all translation activities at each codon, considering both precise and incorrect translations in each sample. By calculating the signal intensity for every codon across the entire genome, we were able to obtain an unbiased estimation of translation error rates for each codon. Notably, owing to the low translation error rate (ranging from 10^−4^ to 10^−3^ per codon site ^1,2^), sampling bias can result in some rare codons (typically unpreferred) not displaying any detected translation errors in the mass spectrometry data. This has the potential to skew estimates of translation error rates significantly and underscores the need for caution when comparing error rates across codons. To address this bias, we employed three strategies, all of which supported the conclusion that optimal codons in *D. melanogaster* exhibit significantly lower translation error rates than their nonoptimal counterparts after correcting for biases. Notably, a recent study in *E. coli* has also presented evidence of higher translational accuracy associated with optimal codons compared to nonoptimal ones ^52^. These findings suggest that codon optimization might represent a conserved strategy across evolution for enhancing translation accuracy in both prokaryotes and eukaryotes.

Initially regarded as neutral ^42,102^, synonymous mutations are now recognized to be subject to either weak ^12,37,43,45^ or strong purifying selection ^44,100,101^. Despite the large effective population size of *D. melanogaster* (estimated at around 10^6^), which should theoretically boost the efficiency of natural selection ^103^, neither previous studies ^13,59-61^ nor this study have found definitive proof of positive selection on synonymous mutations in the lineage of *D. melanogaster* after diverging from *D. simulans*. In contrast, significant signals of positive selection have been detected on the synonymous mutations in the *D. simulans* lineage ^13,59,104^. Notably, a more pronounced mutational bias from C/G toward A/T alleles was inferred in the *D. melanogaster* lineage compared to the *D. simulans* lineage ^62-65,105^. Given that translationally favored codons consistently end with G or C in *Drosophila* ^56^, this mutation bias is anticipated to diminish the prevalence of optimal codons in *D. melanogaster*. Indeed, our examination of codon occurrences in the genomes of *D. melanogaster, D. simulans*, and *D. yakuba* revealed that the prevalence of optimal codons was significantly lower in the *D. melanogaster* genome than in the other two species, or even when compared to the inferred sequences of the most recent common ancestor of *D. melanogaster* and *D. simulans* (Supplementary Fig. 8). The driving force responsible for the shift in mutation bias toward A/T in *D. melanogaster* lineage remains unclear. It is possible that this alteration in mutational bias, in conjunction with changes in effective population size and shifting ecological conditions in *D. melanogaster* after its divergence from *D. simulans*, obscures or diminishes the effectiveness of positive selection on synonymous mutations during evolution in the *D. melanogaster* lineage ^13,59-61^. However, by comparing derived allele frequencies within *D. melanogaster* populations, we have provided support for the hypothesis that natural selection favors codon-optimizing mutations, while codon-deoptimization mutations are prone to negative selection. It’s important to note that not all codon-deoptimizing mutations are necessarily deleterious in *D. melanogaster*, as some may prove advantageous in specific genomic contexts ^106^. In summary, our study effectively bridges the gap between the functional consequences of codon usage bias and the underlying evolutionary driving forces.

Previous studies have revealed the existence of a general trade-off between translation accuracy and speed ^21,22,107^. Such a trade-off has recently been deciphered for the same type of codon occurring at different genomic positions, driven by factors such as magnesium concentration ^23^ and RNA secondary structure ^26^. Our findings, however, propose a different scenario, where optimal codons are both more accurate and faster in translation compared to nonoptimal codons. This suggests that, when examining distinct codons encoding the same amino acid, the conventional accuracy-speed trade-off might not apply. The role of tRNA abundance is pivotal in the translation decoding process, and the competition among tRNA molecules for codon binding impacts translation efficiency and accuracy. Typically, optimal codons are decoded by more prevalent tRNAs, while nonoptimal codons are decoded by rarer tRNAs ^108-114^. It’s plausible that natural selection has influenced tRNA abundance within cells, though further research is necessary to validate this hypothesis.

In addition to regulating translation accuracy and efficiency, synonymous changes may affect precursor mRNA splicing ^115^, alter mRNA secondary structure ^116^, control translation initiation ^117^, and influence co-translational protein folding by modulating local translation elongation rates ^113^. Beyond the harmful effects observed in yeast ^44^ and associations with human diseases ^118,119^, it’s becoming increasingly clear that synonymous mutations may contribute to adaptive evolution more frequently than previously assumed, as seen in microbes ^120^. Our study sheds light on the driving forces behind natural selection on synonymous mutations related to codon optimization and deoptimization. This underscores the significance of gaining a comprehensive understanding of codon usage bias, which in turn enriches our comprehension of the translation process, its influence on cellular functions, and the evolutionary factors shaping codon adaptation.

## Methods

### Identifying translation error events using mass spectrometry data

We downloaded mass spectrometry datasets of 17 distinct samples from 15 representative time points of the Oregon-R strain of *D. melanogaster* from the Proteomics Identifications Database (PRIDE) ^121^ under identifiers PXD005691 and PXD005713 ^70^. These samples encompassed four stages of embryos (0–2h, 4–6h, 10–12h, and 18–20h old), four stages of larvae (L1, L2, early L3, and L3 crawling larvae), five stages of pupae (P1 to P5), virgin males and females at 4 hours after hatching, and one-week-old adult male and female flies. The genome sequencing data of the Oregon-R strain was downloaded from the Sequence Read Archive (SRA) under accession SRP035237 ^122^. We aligned the next-generation sequencing (NGS) reads to the ISO-1 reference genome (FlyBase r6.48, http://www.flybase.org) using Spliced Transcripts Alignment to a Reference (STAR) 2.7.10a ^123^ and identified the single-nucleotide polymorphisms (SNPs) with Freebayes 1.3.5 ^124^. Based on the Oregon-R strain’s genome sequencing results, we generated reference protein sequences for mass spectrometry data processing. MaxQuant 2.1.0.0 ^73^ was used to identify the base and dependent peptides, and translation error events were detected using previously published procedures ^20^. If the same translation error event is identified in more than three samples simultaneously, it will be excluded from subsequent analysis. In addition, potential interference from RNA editing sites is excluded from the identified error translation sites. Potential translation error events on the 217th residue (lysine) of Prm and the 293rd residue (serine) of Adlk2 were excluded from subsequent analyses due to their overlap with known editing sites ^74^.

### Estimation of translation error rates

The coding sequences (CDSs) to which the peptide sequences were aligned were identified based on information parsed from the peptides.txt file in the MaxQuant output data. The mass spectral signal intensities of the dependent peptides were fetched from the matchedFeatures.txt in the MaxQuant output data, following previously published procedures ^20^. For a type of codon *j* (ranging from 1 to 61), we calculated its summed intensity in the proteomic data using the formula

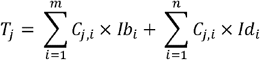

where *C*_*j,i*_ represents the occurrences of the *j*th type of codon in the *i*th peptide, *Ib*_*i*_ and *Id*_*i*_ denote the mass spectral signal intensities of the base or dependent peptides, respectively; *m* and *n* denote the number of base peptides and dependent peptides included in the analysis, respectively.

Similarly, for the *j*th type of codon, we calculated its summed intensity that was erroneously translated using the formula

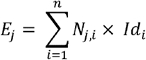

where *N*_*j,i*_ denotes the occurrences of the mistranslated codon *j* in the th-dependent peptides, *N*_*j,i*_ is 1 for all occurrences in this analysis. We then calculated the error rate of the *j*th type of codon as *e*_*j*_ = *E*_*j*_ /*T*_*j*_.

After calculating the translation error rate of each codon, we followed a previous study ^52^ and calculated the relative translation error rate (RMR) of the codons for comparison purposes using the formula

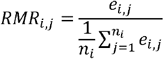

where *e*_*i,j*_ is the average absolute translation error rate of codon *j* of amino acid *i*, and *n*_*i*_ is the number of synonymous codons of amino acid *i*.

### Calculation of RSCU

The gene expression profiles of the developmental time course of *D. melanogaster* determined by the group from Baylor College of Medicine (BCM_1_RNAseq) ^125^ were downloaded from FlyBase (www.flybase.org). For each protein-coding gene, we calculated the median gene expression level (RPKM) across different samples to represent the mRNA expression of that gene. To estimate the codon occurrences used in the *D. melanogaster* transcriptomes, we counted the number of codon occurrences in the longest CDS of each gene. We multiplied that number by the RPKM of the gene. The RSCU value was calculated using the method previously reported ^82,126^, which involves computing the ratio of the number of occurrences of each codon to the average number of occurrences of all codons corresponding to the same amino acid, while considering the usage of each codon in the transcriptome instead of the genome. The coding sequences used for calculations, along with their transcriptomic expression levels and the resulting RSCU values, were documented in Supplementary Data 1.

### Translation error codon characteristic statistics

Based on the identified translation error events (total of 1,374; see Supplementary Data 3; Supplementary Fig. 3), potential numbers of base mismatch occurrences and mismatch positions were inferred (Fig. 1b). Translation errors were primarily assigned to events with the fewest base mismatches. If multiple mismatch occurrences had the same amount of mismatches, the translation error was evenly distributed among them. The calculation of random expectation involved setting the frequency of each codon erroneously translating to other amino acids in Supplementary Data 3 to its codon usage frequency. Translation errors were assigned according to the aforementioned approach. If only one mismatch occurs at a codon position, it is categorized as a near-cognate error; if there is more than one, it is categorized as a noncognate error.

We simulated the relative occurrences of NeCE and NoCE events under randomness by assuming that each codon has an equal probability of being mistranslated into any of the 19 other amino acids, and then calculated the expected numbers of NeCEs and NoCEs events. Taking into account the relative occurrences of each type of codon, we anticipated that 32% of mistranslation events would be NeCEs, and 68% would be NoCE events under random conditions. We then computed the relative frequencies of mismatches at each position of the codon under randomness, resulting in a relative ratio of 5.64: 7.07: 1 for positions 1, 2, and 3 of the codon, respectively. In contrast, among the 788 observed NeCE events, the relative ratio was 8.00: 4.85: 1 for positions 1, 2, and 3 of the codon, respectively. Therefore, compared to random expectations, the observed NeCE events exhibited a substantial enrichment in the first position (8.00/5.64 = 1.42) and a depletion in the second position (4.85/7.07 = 0.686) relative to the third position of the codon (Fig. 1e).

### Simulating the effect of mass spectrometry throughput on translation error detection

The translation error rate for all codons was set to 2.37×10^−4^ errors per amino acid. In each run of the simulation, a total number of amino acids detected was assigned to the codons based on their occurrences in the transcriptomes of *D. melanogaster*. The number of translation error events on a codon was assumed to follow a Poisson distribution. Each simulation run included a range of amino acid numbers in the proteomes from 5×10^4^ to 5×10^6^, with an interval of 5×10^4^. The simulation was independently repeated 100 times.

### Linear mixed models

The linear mixed models were fitted using the “mixedlm” function from the Python package Statsmodels (<u>pypi.org/project/statsmodels</u>; v.0.13.5) to examine the relationship between translation error and codon usage (RSCU).

### Calculation of ribosome density distribution

We reanalyzed the mRNA-Seq and Ribo-Seq data we had previously generated for different stages of the ISO-1 strain of *D. melanogaster* (Sequence Read Archive (SRA) accession number SRP067542) ^71^. After eliminating tRNA- and rRNA-related reads, raw reads were aligned to the reference genome of *D. melanogaster* using STAR 2.7.10a ^123^. The corresponding sites of the ribosome A-site on different length (25–35 nt) reads were estimated using plastid 0.5.1 ^127^, and the A-site occupancy values corresponding to each site on the transcript with the longest CDS in the gene were counted. Because regions at the beginning and end of the CDS can cause abnormal ribosome blockage due to translation initiation or termination, we omitted the first 15 and last 5 codons of the CDS. We eliminated genes with insufficient RPF coverage (RPF coverage < 0.5) and short coding regions (< 150 codons). To exclude the effect of different translation initiation rates between genes on ribosome density, the ribosome density on each codon was normalized by dividing it by the gene’s average codon ribosome density.

### Inference of the common ancestral sequences of *D. melanogaster* and *D. simulans*

FASTA alignments of 26 insects with *D. melanogaster* for CDS regions were downloaded from the UCSC Genome Browser (genome.ucsc.edu). Using *D. yakuba* as an outgroup, the common ancestral CDS sequences of *D. melanogaster* and *D. simulans* were inferred. If the nucleotide sequences at a certain position were not the same among all three species, the ancestral sequence of that position was labeled as unclear, and the codons associated with that position were excluded in subsequent calculations.

### McDonald-Kreitman test for synonymous mutations in the *D. melanogaster* populations

The genome sequence alignments between *D. melanogaster, D. simulans*, and *D. yakuba* were retrieved from the UCSC Genome Browser (genome.ucsc.edu). We inferred the ancestral states of the synonymous mutations that are polymorphic in the *Drosophila* Genetic Reference Panel 2 (DGRP2) database ^89,90^ of *D. melanogaster* by comparing the nucleotides at the orthologous sites of *D. melanogaster, D. simulans*, and *D. yakuba*. The synonymous mutations were separated into two categories, optimization and deoptimization, based on whether the codon RSCU value increased or decreased after mutation. Mutations in the 8–30 nt region of the short introns (≤65 nt) were parsed as previously described ^71^. After correcting for multiple substitutions of fixed sites between *D. melanogaster* and *D. simulans* using Kimura’s 2-Parameterm model ^128^, we calculated the α values of synonymous mutations using the AsymptoticMK method ^83,84^. Mutations with minor allele frequency < 0.05 were excluded in the AsymptoticMK tests.

We classified *D. melanogaster* genes into three groups based on their median expression levels across samples: highly expressed (4,565 genes), moderately expressed (4,817 genes), and lowly expressed (4,517 genes). We calculated the α values of synonymous mutations in each class of genes using the same neutral controls in the AsymptoticMK tests.

## Acknowledgments

We thank Dr. Mengyi Sun for the constructive comments on the manuscript. This work was supported by grants from the Ministry of Science and Technology of the People’s Republic of China (2022YFE0132000), the National Natural Science Foundation of China (32070597, 32122022), the Natural Science Foundation of Beijing (5212006), and the SLS-Qidong Innovation Fund. Some analyses were performed on the High-Performance Computing Platform of the Center for Life Sciences. We thank the National Center for Protein Sciences at Peking University for technical assistance.

