## Supplemental Material for "Genome-wide impact of codon usage bias on translation optimization in *Drosophila melanogaster*"

**FIGURE and TABLE LEGEND**

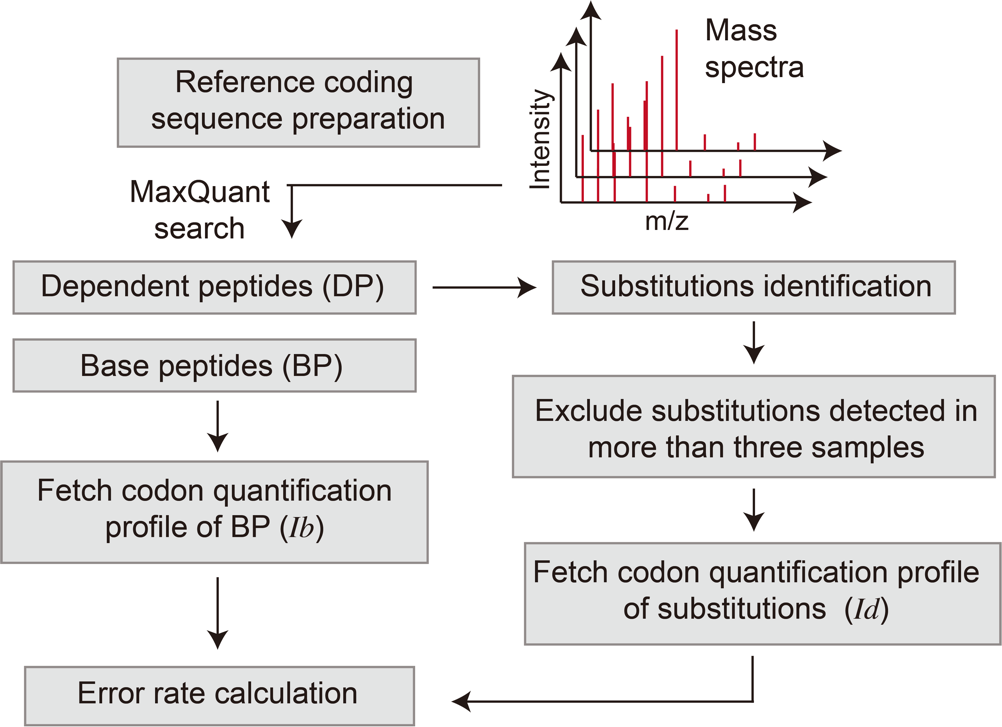

**Supplementary Fig. 1. The workflow for identifying translation error events and** **estimating translation error rates at the codon level.**

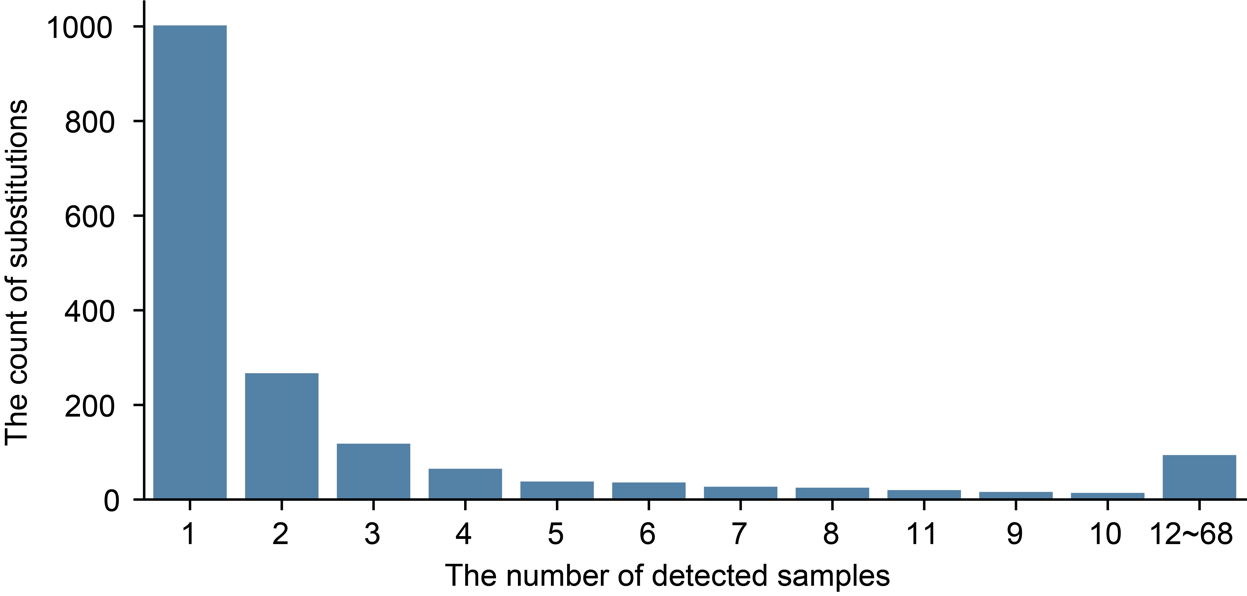

**Supplementary Fig. 2. The number of samples (out of 68 samples) in which a candidate translation error occurrence was identified.**

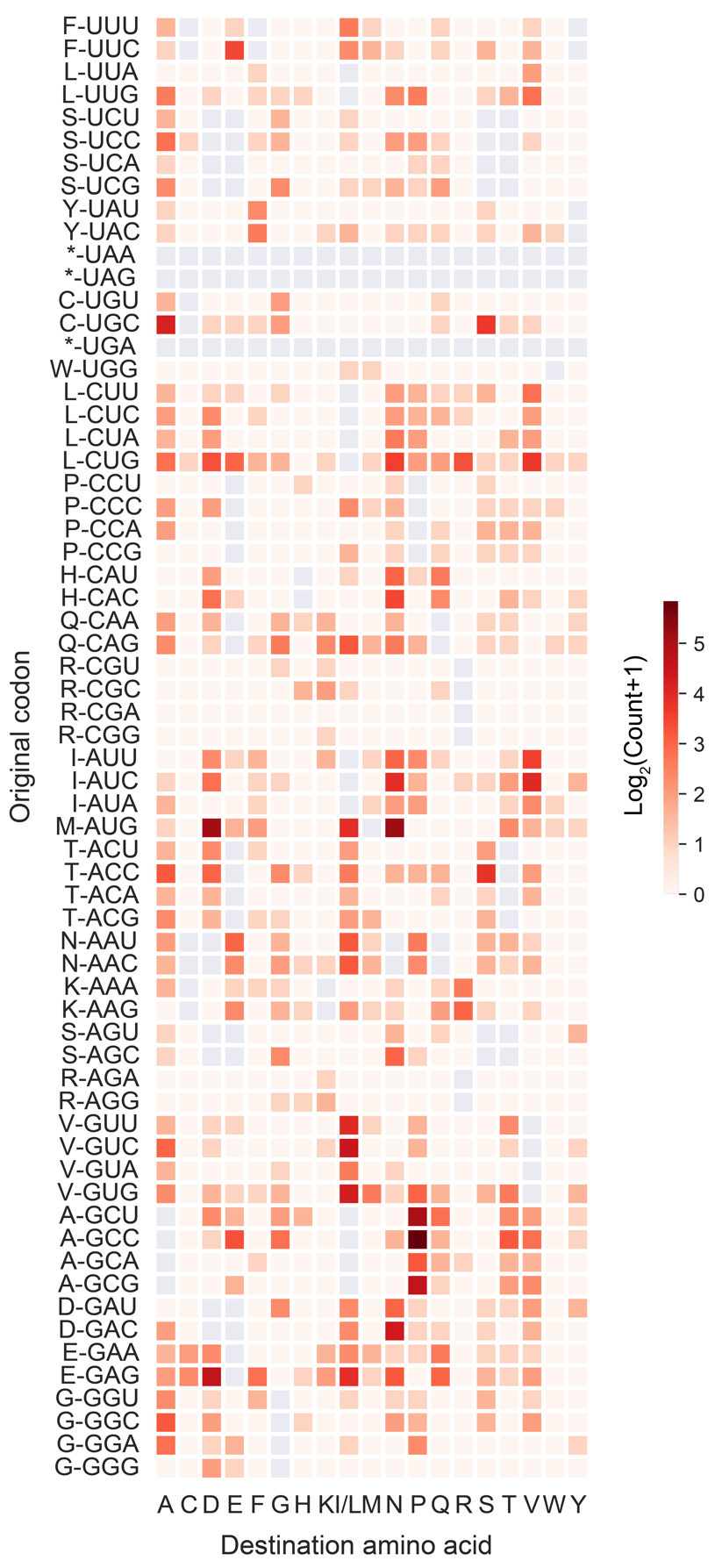

**Supplementary Fig. 3. Overview of detected amino acid substitution events at 1,374 distinct genomic sites across the 68 libraries.** The vertical axis is the original codon on the CDS, and the horizontal axis is the amino acid after substitution. The shade of the color represents the number of genomic sites that showed a particular mistranslation event, and the gray color means that the substitution event does not exist or could not be confirmed by means of mass difference.

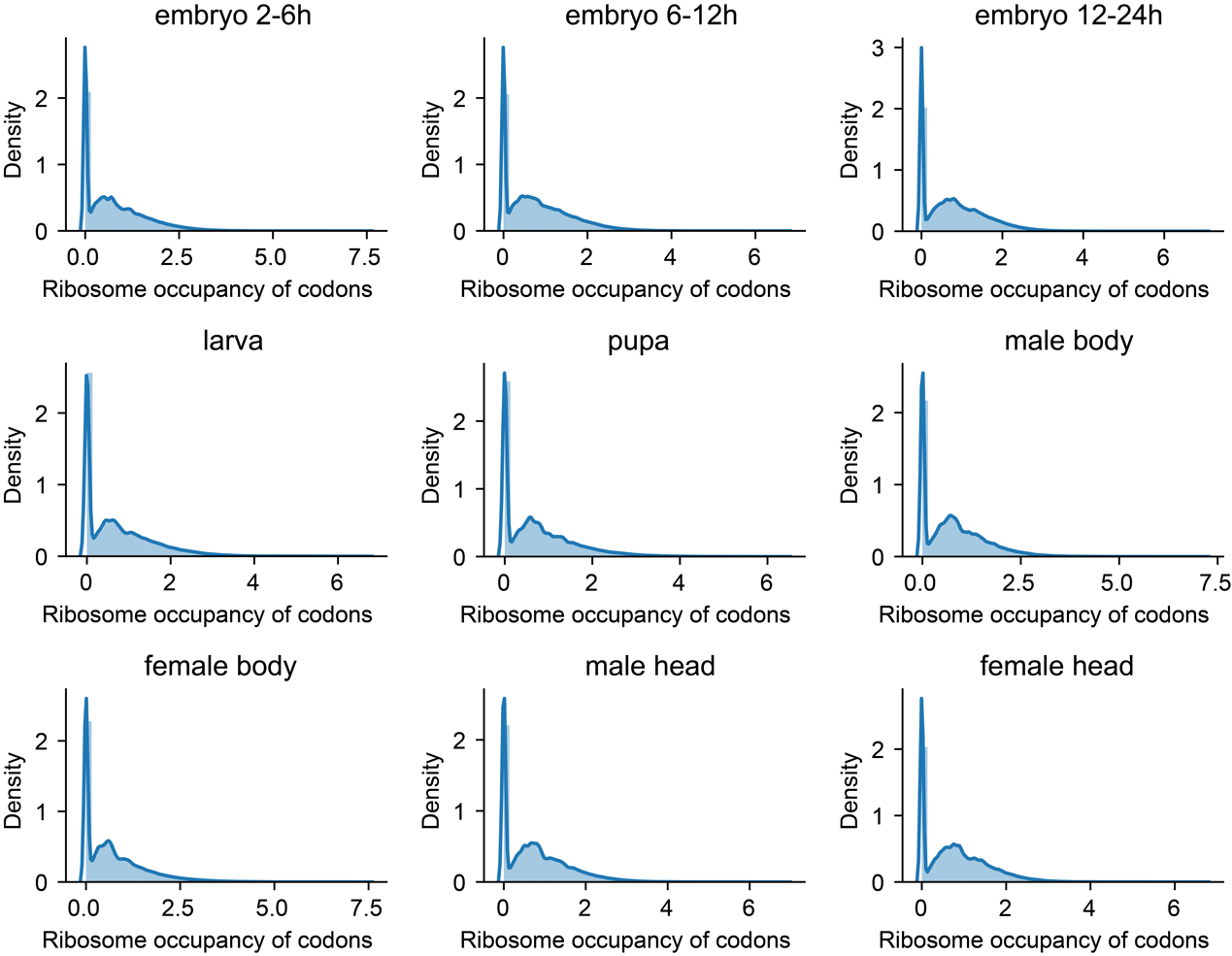

**Supplementary Fig. 4. The distribution of log2-transformed normalized A-site occupancy of different samples of *D. melanogaster*** (zero-RPF covered codons were retained).

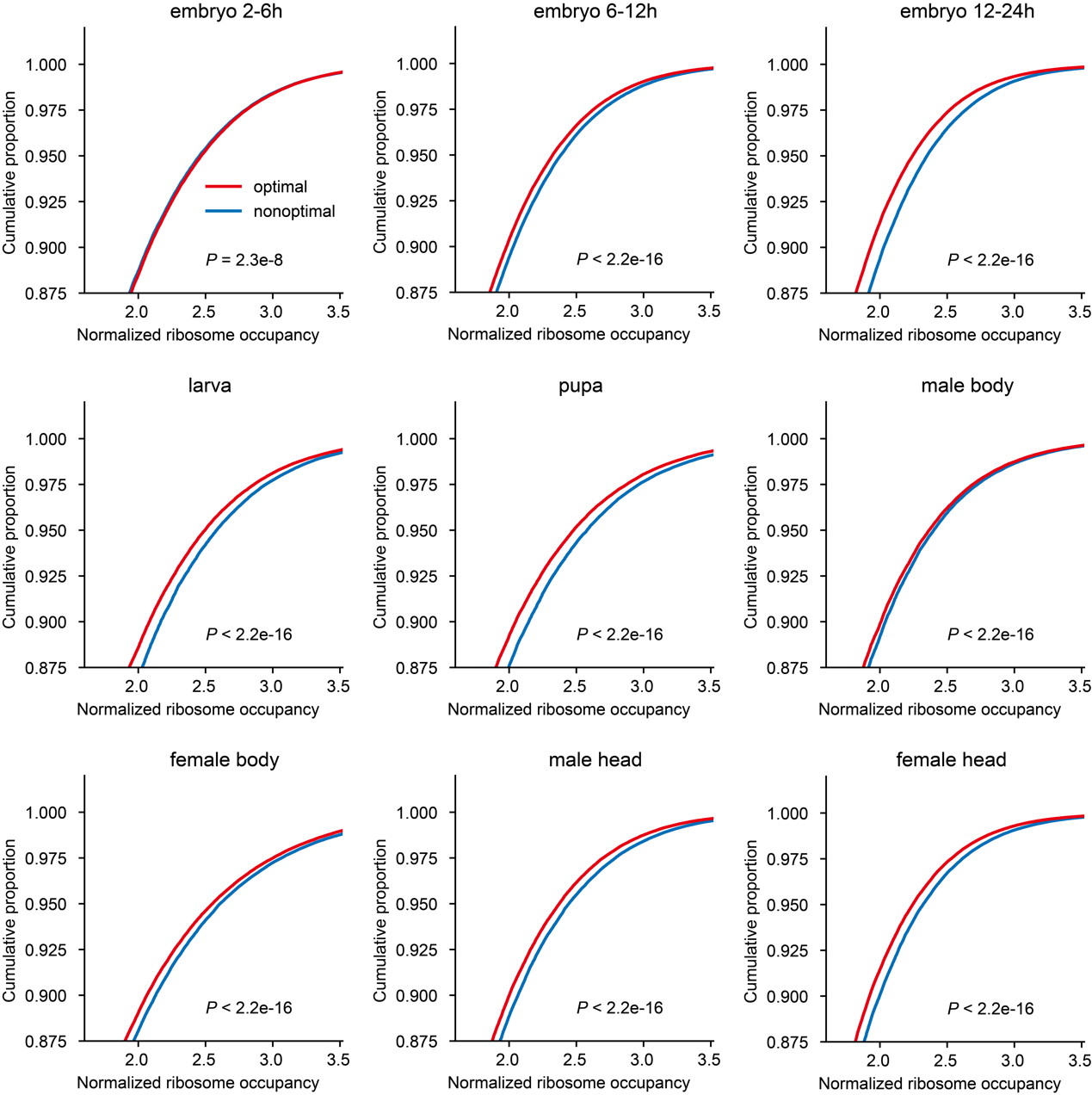

**Supplementary Fig. 5. Comparison of normalized ribosome densities between optimal and nonoptimal codons in other samples of *D. melanogaster*.** *P* value was calculated using the Wilcoxon rank-sum test.

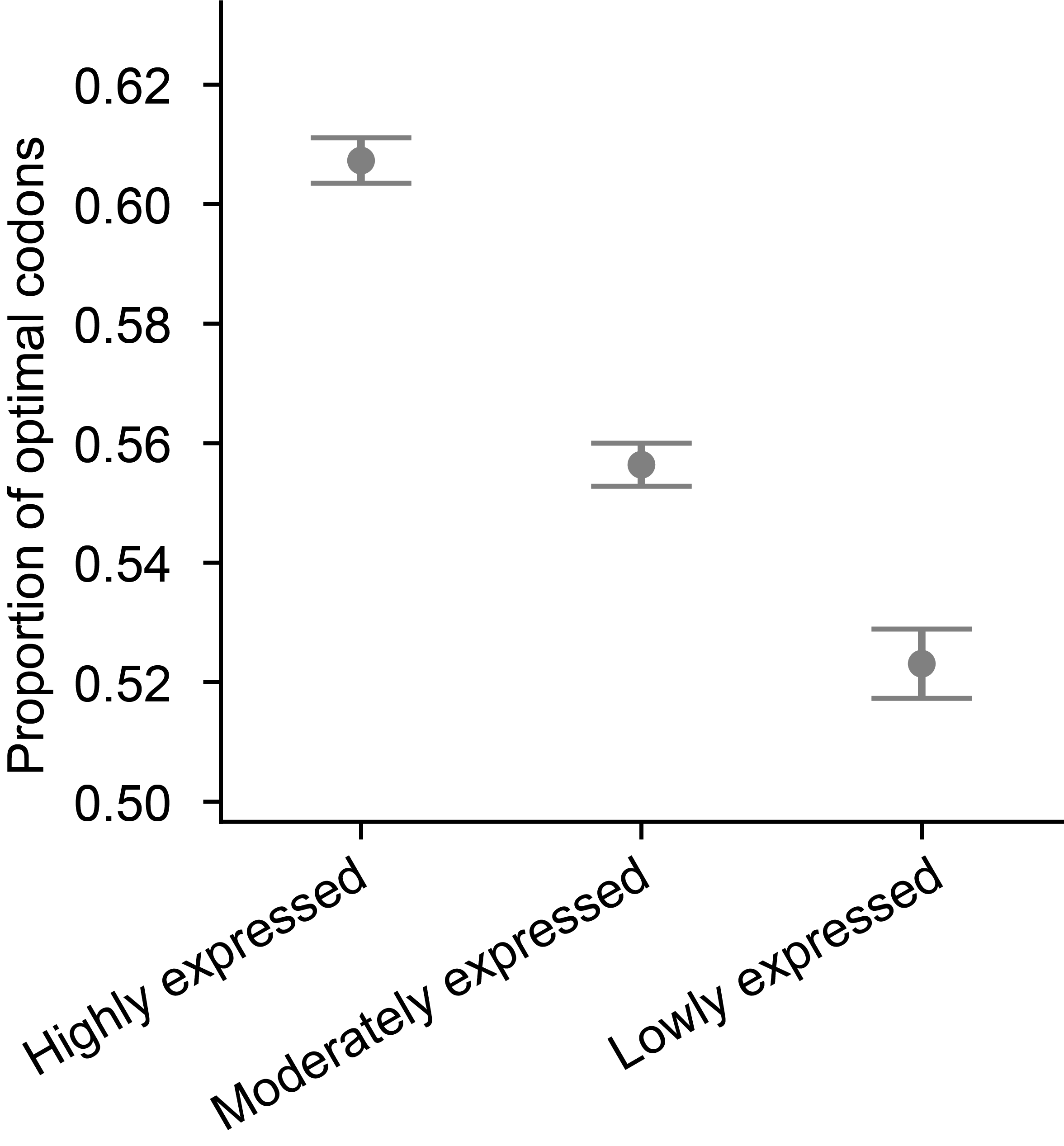

**Supplementary Fig. 6. The proportion of optimal codons among the three classes of genes with different expression levels.** The bootstrap method was used to estimate the mean values and 95% confidence intervals.

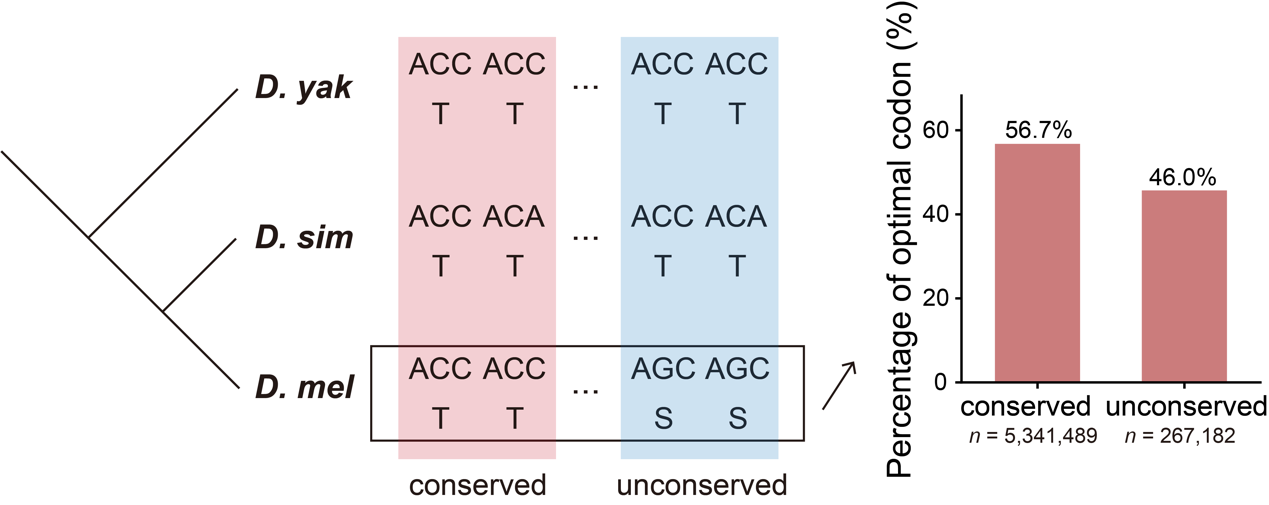

**Supplementary Fig. 7. A significantly higher percentage of optimal codons are observed in conserved amino acid sites across species than in nonconserved sites.** The analysis retained only the amino acids that were identical in both *D. yakuba* and *D. simulans*. Among them, the conserved category refers to positions where the amino acids are the same between *D. simulans* and *D. melanogaster*, while the unserved category refers to positions where there are differences in amino acids between *D. simulans* and *D. melanogaster*.

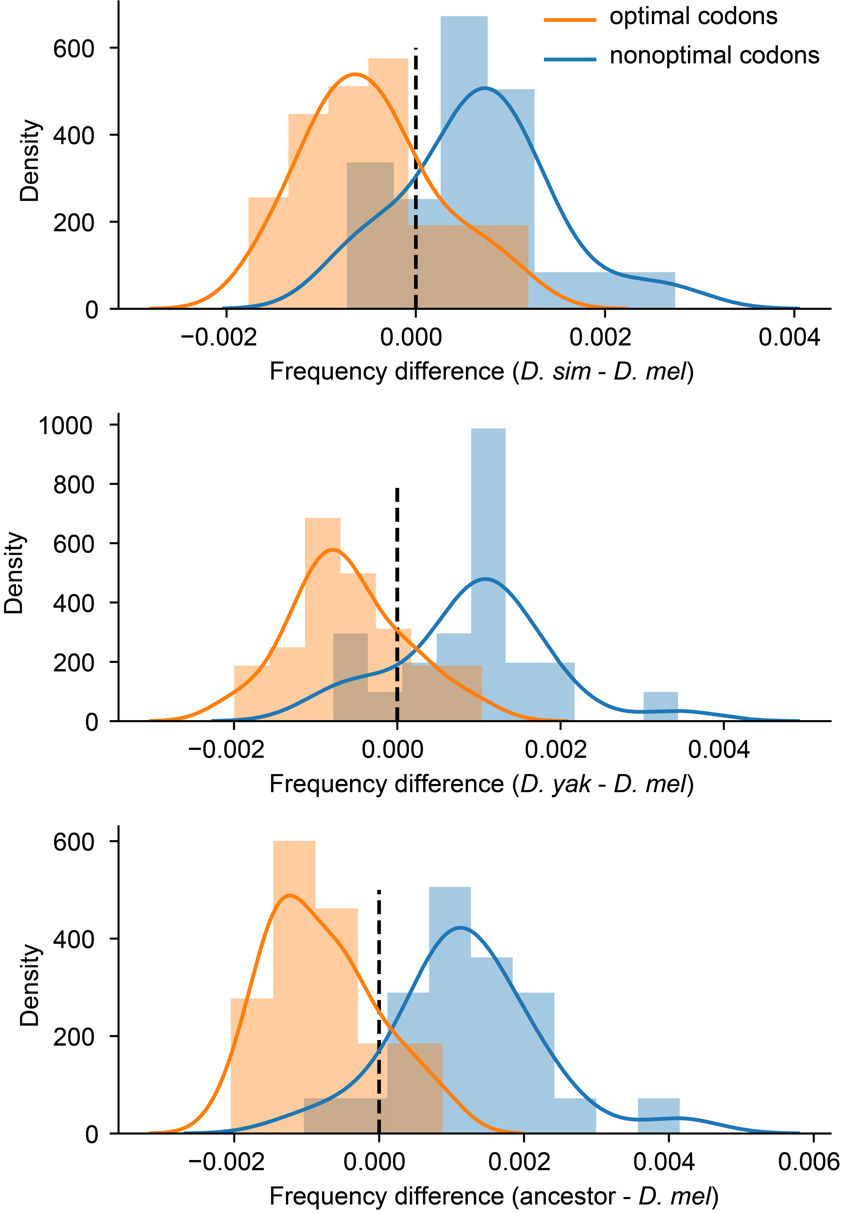

**Supplementary Fig. 8. Differences in codon usage frequency across different species.** The *x*-axis shows the difference in codon usage frequency, comparing specific codons between *D. simulas* and *D. melanogaster* (top panel), between *D. yakuka* and *D. melanogaster* (middle panel), and in the most recent common ancestor of *D. simulans* and *D. melanogaster*, relative to *D. melanogaster* itself (bottom panel).The *y*-axis represents the distribution density of the corresponding codons. Optimal codons are shown in orange, while nonoptimal codons are shown in blue.

**Supplementary Data 1. Codon usage** **preferences in *Drosophila melanogaster*.** The data includes the RSCU values for each codon, as well as the mean expression level (RPKM) and coding sequences of genes used for calculating codon usage.

**Supplementary Data 2. The mistranslation rate for each of the 61 codons in the 68 mass spectrometry samples.**

**Supplementary Data 3. Statistics of specific types of codon (****origin) translation errors into amino acids (****destination).** In total 1,374 translation error events were identified.
